# Meta-Analysis of Transcriptomic Datasets Reveals Key Immune Gene Profiles and Signaling Pathways in Bos taurus

**DOI:** 10.1101/2025.09.13.676061

**Authors:** Vennila Kanchana Devi Marimuthu, Kishore Matheswaran, Menaka Thambiraja, Ragothaman M Yennamalli

## Abstract

Understanding immune mechanisms in cattle is vital for improving disease resistance through informed breeding and vaccine development. Meta-analysis enables the integration of multiple transcriptomic studies to identify consistent gene expression patterns and enhance statistical power. In this study, we performed a meta-analysis of four bovine transcriptomic datasets (GSE45439, GSE62048, GSE125964, and GSE247921) to identify immune-related differentially expressed genes (DEGs) in *Bos taurus*. These datasets covered infections with *Mycobacterium bovis* and *Mycobacterium avium* subsp. *paratuberculosis*, comparing diseased and healthy cattle. Our pipeline included FastQC, Trimmomatic, Bowtie2, SAMtools, FeatureCounts, DESeq2, and MetaRNASeq, leading to the discovery of 28 significant DEGs (12 upregulated and 16 downregulated). Comparison with an innate immune gene database identified five key immune-related genes—IL1A, RGS2, RCAN1, and ZBP1—with known regulatory roles in immunity. KEGG enrichment analysis revealed involvement in Necroptosis, Osteoclast Differentiation, Oxytocin Signaling, and cGMP–PKG Signaling pathways, associated with inflammatory cell death, cytokine signaling, and immune cell differentiation. This meta-analysis enhances understanding of conserved immune signaling mechanisms in cattle and highlights genes that may serve as biomarkers for immune competence, disease susceptibility, and vaccine responsiveness, offering valuable insights for future bovine immunogenomics research.

## 1. Introduction

Cattle represent one of the most economically significant livestock species globally, contributing substantially to food security, agricultural economies, and livelihoods across diverse geographical regions□(World Bank, 2023). However, infectious diseases continue to pose major threats to cattle health and productivity, resulting in substantial economic losses estimated in billions of dollars annually (Drake, 2021). Diseases such as tuberculosis caused by *Mycobacterium bovis*, Johne’s disease caused by *Mycobacterium avium* subspecies *paratuberculosis* (MAP), and various other infectious agents significantly impact bovine welfare and agricultural sustainability□(Thoen et al., 2019, Mullie et al., 2021). The emergence of antibiotic resistance and growing concerns about antimicrobial residues in food products have intensified the need for alternative approaches to disease management in cattle (World Health Organization, 2017).

In India, livestock management faces unique challenges that significantly amplify the impact of infectious diseases on cattle populations. The prevalence of bovine tuberculosis (bTB) among farm and dairy cattle in India is estimated to be around 7.3%, which makes it a country with one of the largest infected herds in the world (Srinivasan et al., 2018). This translates to an estimated 21.8 million infected cattle in India a population greater than the total number of dairy cattle in most developed countries (Ramanujam & Palaniyandi, 2023). The situation is particularly concerning in peri-urban smallholder dairy farms, where intensive, industry-style livestock rearing has led to emerging vulnerabilities at the human- animal-environment interface (Chauhan et al., 2019). Unlike developed countries where systematic control programs have successfully reduced disease prevalence, India’s livestock sector continues to grapple with limited diagnostic infrastructure, inadequate surveillance systems, and challenges in implementing comprehensive disease control measures. The economic implications are substantial, as India’s dairy industry the world’s largest milk producer (Food and Agriculture Organization of the United Nations, 2025) relies heavily on smallholder farmers who often lack the resources for proper disease management and prevention strategies.

Understanding the molecular mechanisms underlying immune responses in cattle has become increasingly crucial for developing effective disease prevention and treatment strategies. The bovine immune system, while sharing fundamental similarities with other mammalian species, exhibits unique characteristics that require species-specific investigation (Vlasova et al., 2021). Genetic variation in immune competence among cattle breeds and individuals presents both challenges and opportunities for improving disease resistance through selective breeding programs (Verma et al., 2016). The identification of key immune genes and pathways could facilitate the development of genetic markers for disease susceptibility, inform breeding decisions, and guide the design of more effective vaccines and therapeutic interventions.

Traditional approaches to studying bovine immunity have relied heavily on phenotypic observations, serological assays, and candidate gene approaches (Mallikarjunappa et al., 2022) However, these methods often provide limited insights into the complex molecular networks governing immune responses. The advent of high-throughput genomic technologies, particularly RNA sequencing (RNA-Seq), has revolutionized our ability to comprehensively characterize gene expression patterns and identify molecular signatures associated with immune activation, pathogen recognition, and disease susceptibility in cattle (Byron et al., 2016).

Earlier transcriptomic studies have advanced our understanding of bovine immune responses to infection and inflammation. These prior investigations emphasize both the complexity of host pathogen interactions and the challenges of deriving generalizable conclusions from isolated experimental systems. For example, Nalpas et al. (2013) (GSE45439) analyzed the transcriptional responses of bovine monocyte derived macrophages (MDMs) to *Mycobacterium bovis* infection using strand-specific RNA sequencing (Nalpas et al., 2013). Bovine tuberculosis is a chronic infectious disease in cattle caused by the bacterium *Mycobacterium bovis*, a significant zoonotic pathogen transmitted primarily through inhalation of infected droplets or contact with contaminated bodily fluids. In cattle, it can cause chronic wasting, respiratory issues, and enlarged lymph nodes. Their study identified 2,584 differentially expressed (DE) genes (1,392 upregulated; 1,192 downregulated) on the sense strand and 757 on the antisense strand, with prominent involvement in innate immune pathways, apoptosis, and cell signaling. Genes such as IL17A, SAA3, ADRB3 and M-SAA3.2 were strongly upregulated and GABBR2, CDH26, SLC16A12, FAIM2 and SH3TC2 were strongly downregulated indicating activation of macrophage mediated inflammatory responses. Notably, antisense transcripts also played a role in modulating gene expressions, offering insights into post transcriptional regulatory mechanisms (Nalpas et al., 2013).

Casey et al. (2015) (GSE62048) examined early immune responses to *Mycobacterium avium* subspecies *paratuberculosis* (MAP), the causative agent of Johne’s disease, in bovine MDMs (Casey et al., 2015). Johne’s disease is a chronic intestinal infection in ruminants caused by MAP, leading to weight loss, diarrhea, reduced milk production, and eventual death. It is primarily transmitted through the fecal-oral route and poses significant economic and animal health concerns in the cattle industry. Their study identified 245 differentially expressed (DE) genes at 2 hours post-infection (209 upregulated; 36 downregulated) and 574 DE genes at 6 hours post-infection (342 upregulated; 232 downregulated). Key upregulated genes included CSF3, CXCL3, TNFAIP6, CCL20, and IL1B at 2 hpi, and LOXL4, GJB2, FFAR4, STOML3, and SAA3 at 6 hpi. Notable downregulated genes were RAB3A, OSM, FOS, POU3F1, and ANKRD63 at 2 hpi, and TNFSF18, KIT, SLC7A8, STON2, and ARHGAP26 at 6 hpi. These genes were involved in IL-10 signaling, TLR2/4, and NOD2-mediated recognition pathways, indicating a dynamic and time-dependent macrophage immune response to MAP infection. The study highlighted the enhanced sensitivity of RNA-Seq over microarrays in detecting early immune gene perturbations.

Similarly, Brewer et al. (2020) (GSE125964) analyzed endometrial gene expression in cows seven days postpartum (DPP) to identify predictive signatures of uterine disease (Brewer et al., 2020). Postpartum uterine diseases in cattle are a significant economic concern due to their impact on fertility. These conditions, primarily caused by bacterial contamination of the uterus after calving, can lead to inflammation, delayed uterine involution, and disruptions in ovarian function, ultimately affecting conception rates. Clinical endometritis is characterized by purulent or mucopurulent uterine discharge occurring after 21 days postpartum, while subclinical endometritis is defined by a high proportion of neutrophils in uterine cytology samples collected between 21 and 47 days postpartum. The uterine lumen is particularly susceptible to bacterial contamination during and after parturition. Cows that developed purulent vaginal discharge (PVD) exhibited 297 DE genes (61 upregulated; 236 downregulated), with significant upregulation of IL1A, IL1B, IL17F, and CXCL8, highlighting a heightened inflammatory response. In contrast, only three genes were differentially expressed genes (two upregulated; one downregulated) in cows that developed cytological endometritis (CYTO), suggesting subtler molecular changes (Brewer et al., 2020). Protective immune related genes such as NTS, CCL28, and lactoferrin were enriched in healthy animals, pointing to their potential role in uterine homeostasis.

Recently, Fong et al. (GSE247921) explored the molecular mechanisms of Johne’s disease by analyzing the effects of MAP infection in a bovine mammary epithelial cell line (MAC-T), with and without functional interleukin-10 receptor alpha (IL10Rα) (Fong et al., 2023). Using CRISPR/Cas9-mediated knockout of IL10Rα, the authors compared transcriptional profiles across four experimental groups (wild-type and knockout, with and without MAP infection). A total of 1,388 DE genes were identified between WT and WT-MAP, 1,738 between WT and KO, 1,613 between WT-MAP and KO-MAP, and 561 between KO and KO-MAP, highlighting distinct transcriptional changes across all experimental conditions. The immune response differed between wild-type and IL10Rα knockout MAC-T cells, with minimal response observed between infected and uninfected knockout cells, highlighting IL10Rα’s role in MAP pathogenesis. The study also identified genes involved in chemokine/cytokine signaling, interleukin, and toll-like receptor pathways. These findings support IL10Rα’s importance in coordinating the immune response to MAP infection.

While these studies provide valuable but varied insights into bovine immunogenomics across different tissues and infection models, the variations in experimental design, infection type, timepoints, and tissue specificity contribute to diverse gene level findings. These limitations underscore the need for robust meta-analytical approach to extract conserved transcriptional patterns and elucidate core immune mechanisms underlying bovine disease resistance. To address the critical need for a comprehensive understanding of bovine immune gene expression patterns and to overcome the limitations of individual studies, we conducted a systematic meta-analysis of bovine transcriptomic datasets focusing on immune-related conditions and pathogen infections. Our primary objective was to identify consistently differentially expressed genes across multiple independent studies and to characterize the biological pathways and molecular mechanisms underlying bovine immune responses.

We selected four high quality RNA-Seq datasets from the NCBI Gene Expression Omnibus (GEO) database, representing diverse immune challenges in *Bos taurus*. These datasets encompassed infections with *Mycobacterium bovis* and *Mycobacterium avium* subspecies *paratuberculosis*, as well as clinical and subclinical uterine diseases, providing a comprehensive representation of both infectious and inflammatory conditions affecting cattle health and productivity. The selection criteria emphasized studies with well-defined experimental designs, adequate sample sizes, and clear comparisons between diseased and healthy control groups.

This comprehensive meta-analytical approach is expected to provide several key advantages over individual study analyses. First, by integrating multiple datasets, we can achieve greater statistical power for detecting genes with consistent but modest effect sizes that might be missed in smaller individual studies. Second, the meta-analysis can help distinguish genuine biological signals from study-specific technical artifacts or experimental noise. Third, the identification of genes that are consistently differentially expressed across diverse immune challenges may reveal fundamental mechanisms of bovine immune function that are broadly relevant across different pathological conditions.

The goal of this research is to identify key immune genes and pathways that are consistently activated or suppressed in response to immune challenges from diverse conditions. Expanding the workflow with more datasets would enable the development of genetic selection strategies for disease resistance, guide the design of more effective vaccines, and contribute to the overall improvement of cattle health and welfare.

## 2. Results

The volcano plot generated from the meta-analysis represents the distribution of genes based on their mean log fold change and the log of the combined p-values for gene expression (Figure 1). Here, genes identified as statistically significant are marked in red, while those deemed non-significant appear in grey. Among the 199 common genes identified, 28 significant genes were selected based on a fold change ≥ ±1.5 and p < 0.05 (Saravanan et al., 2021). From this subset, comparison with InnateDB innate immunity genes and immune-related gene key files revealed eight immune genes: IL1A, RGS2, RCAN1, ZBP1, TIMD4, PPARG, SPAG5, and TLR10. Noteworthy genes such as LIPK, RGS2, and MYCBPAP stand out due to their substantial fold change (−13.18, −7.86, and +6.67, respectively) and p-value (1.81×10^−1^□^3^, 5.37×10^−3^□, and 1.58×10^−^□□). Among the immune genes, RGS2, RCAN1, and IL1A notably stand out with both large fold change of −7.86, +3.57, and −3.11 and highly significant p-values (5.37×10^−3^□, 3.63×10^−11^, and 1.36×10^−1^□). This plot enables the clear identification of differentially expressed genes, offering valuable insights into their potential roles in the immune response mechanisms studied in the dataset.

**Figure 1:**
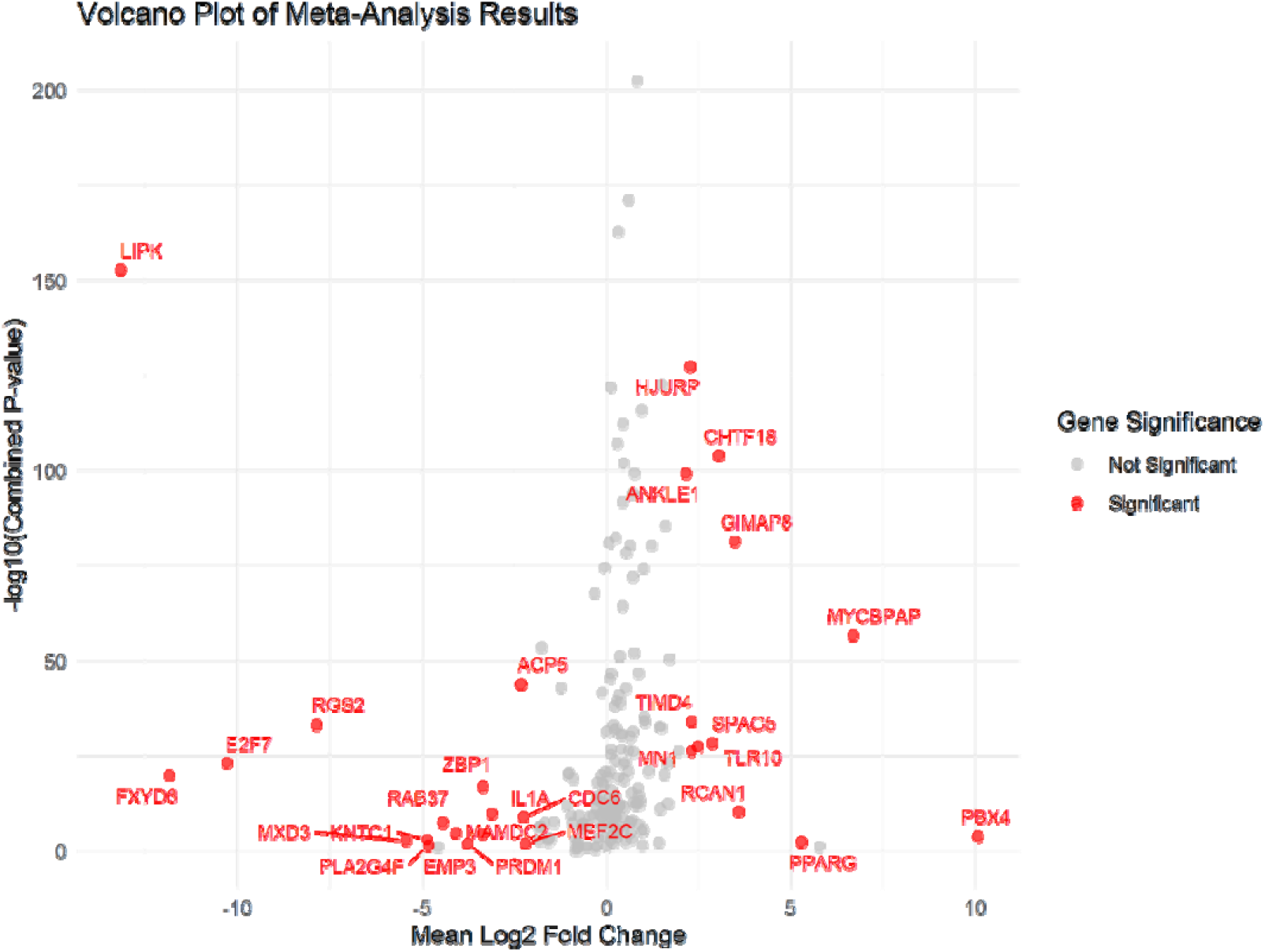
Volcano plot of Significant Genes. The Volcano plot shows the significantly Differentially Expressed Genes (DEG’s) based on fold change and combined p-values from meta-analysis. The X-axis captures the average log fold change, thereby indicating upregulation with positive values and downregulation with negative values. The y-axis reflects statistical significance, where larger values denote stronger evidence against the null hypothesis.

### 2.1. Differential Expression of Genes

Transcriptomic profiling provides a powerful lens to uncover genes that are differentially expressed in response to pathogenic challenges and in understanding the genetic basis of immune responses in cattle. Our Meta-analysis of multiple RNA-Seq datasets show genes that are differentially expressed across all datasets (Figure 2 and Table 1).

**Table 1:** Differentially Expressed Genes Identified Through Meta-Analysis. Significantly DEG’s identified from a meta-analysis of *Bos taurus* RNA-Seq datasets. Immune-related genes were annotated using InnateDB.

| Gene | Combined Pvalue | Mean_Log2FC | Immune Gene |
| --- | --- | --- | --- |
| **Upregulated Genes:** |  |  |  |
| PBX4 | 0.000162 | 10.0574 | NO |
| MYCBPAP | $158 \times 10^{-57}$ | 6.67381 | NO |
| PPARG | 0.00477 | 5.27563 | YES |
| RCAN1 | $363 \times 10^{-11}$ | 3.57311 | YES |
| GIMAP8 | $337 \times 10^{-82}$ | 3.48134 | NO |
| CHTF18 | $11 \times 10^{-104}$ | 3.03795 | NO |
| TLR10 | $406 \times 10^{-29}$ | 2.8542 | **YES** |
| SPAG5 | $383 \times 10^{-28}$ | 2.46975 | **YES** |
| MN1 | $471 \times 10^{-27}$ | 2.31087 | NO |
| TIMD4 | $87 \times 10^{-35}$ | 2.30529 | **YES** |
| HJURP | $39 \times 10^{-128}$ | 2.25407 | NO |
| ANKLE1 | $51 \times 10^{-100}$ | 2.15654 | NO |
| **Downregulated Genes:** |  |  |  |
| MEF2C | 0.011604 | -2.2004 | NO |
| CDC6 | $115 \times 10^{-9}$ | -2.2653 | NO |
| ACP5 | $117 \times 10^{-44}$ | -2.3252 | NO |
| IL1A | $136 \times 10^{-10}$ | -3.1098 | **YES** |
| ZBP1 | $117 \times 10^{-17}$ | -3.3592 | **YES** |
| MAMDC2 | $293 \times 10^{-5}$ | -3.3602 | NO |
| PRDM1 | 0.013848 | -3.7614 | NO |
| EMP3 | $166 \times 10^{-5}$ | -4.0873 | NO |
| RAB37 | $252 \times 10^{-8}$ | -4.4346 | NO |
| PLA2G4F | 0.04345 | -4.8211 | NO |
| KNTC1 | 0.001363 | -4.8693 | NO |
| MXD3 | 0.002161 | -5.4329 | NO |
| RGS2 | $537 \times 10^{-34}$ | -7.8615 | **YES** |
| E2F7 | $778 \times 10^{-24}$ | -10.295 | NO |
| FXYD6 | $151 \times 10^{-20}$ | -11.863 | NO |
| LIPK | $18 \times 10^{-153}$ | -13.18 | NO |

**Table 2:** The data set used in this study. The datasets were selected specifically for those that had comparisons between control and infected samples.

| GENE EXPRESSION<br>OMNIBUS (GEO)<br>SERIES ID | ACCESSION ID | ORGANISM | CONDITION |
| --- | --- | --- | --- |
| GSE45439 | PRJNA194043 | *Bos taurus* | *Mycobacterium bovis* infected<br>vs Control uninfected |
| GSE62048 | PRJNA263027 | *Bos taurus* | Control vs *Mycobacterium*<br>*avium* subsp. paratuberculosis,<br>(MAP) Infected |
| GSE125964 | PRJNA518101 | *Bos taurus* | Healthy vs Clinical & Sub<br>Clinical Postpartum Uterine<br>disease |
| GSE247921 | PRJNA1040910 | *Bos taurus* | *Mycobacterium avium* subsp.<br>paratuberculosis, (MAP)<br>infection vs No treatment |

**Figure 2:**
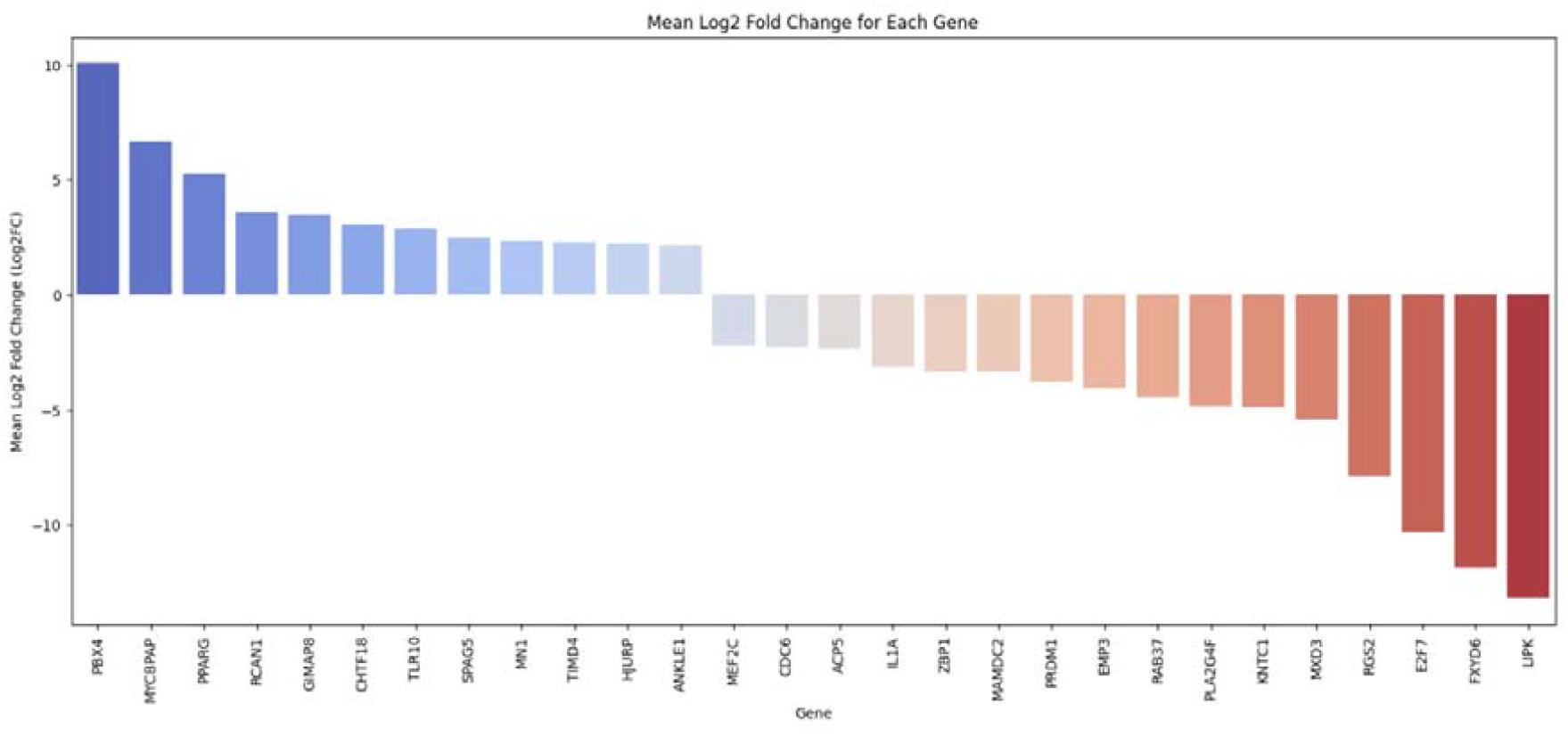
Upregulated and Downregulated Genes found statistically significant among all the datasets. The bar plot shows the significantly upregulated and downregulated genes identified by mean log2 fold change from the meta-analysis.

Figure 3 shows the distribution of mean Log2 fold change values for differentially expressed genes (DEGs) identified in the *Bos taurus* RNA-Seq meta-analysis and shows a balance between upregulated and downregulated genes.

**Figure 3:**
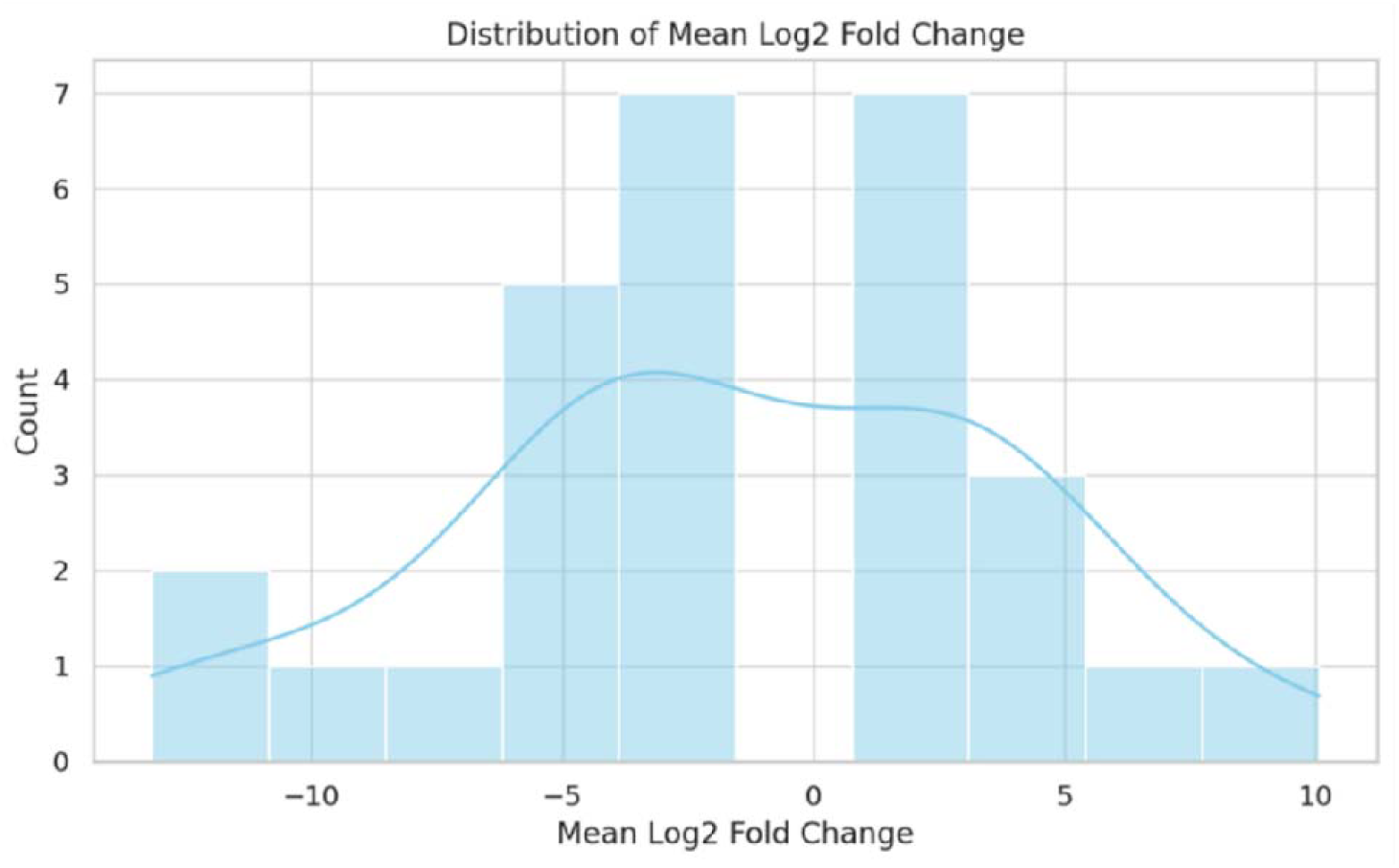
Distribution of Mean Log2Fold Change. The histogram displays the distribution of mean Log2 fold change values for differentially expressed genes across datasets.

**Figure 4:**
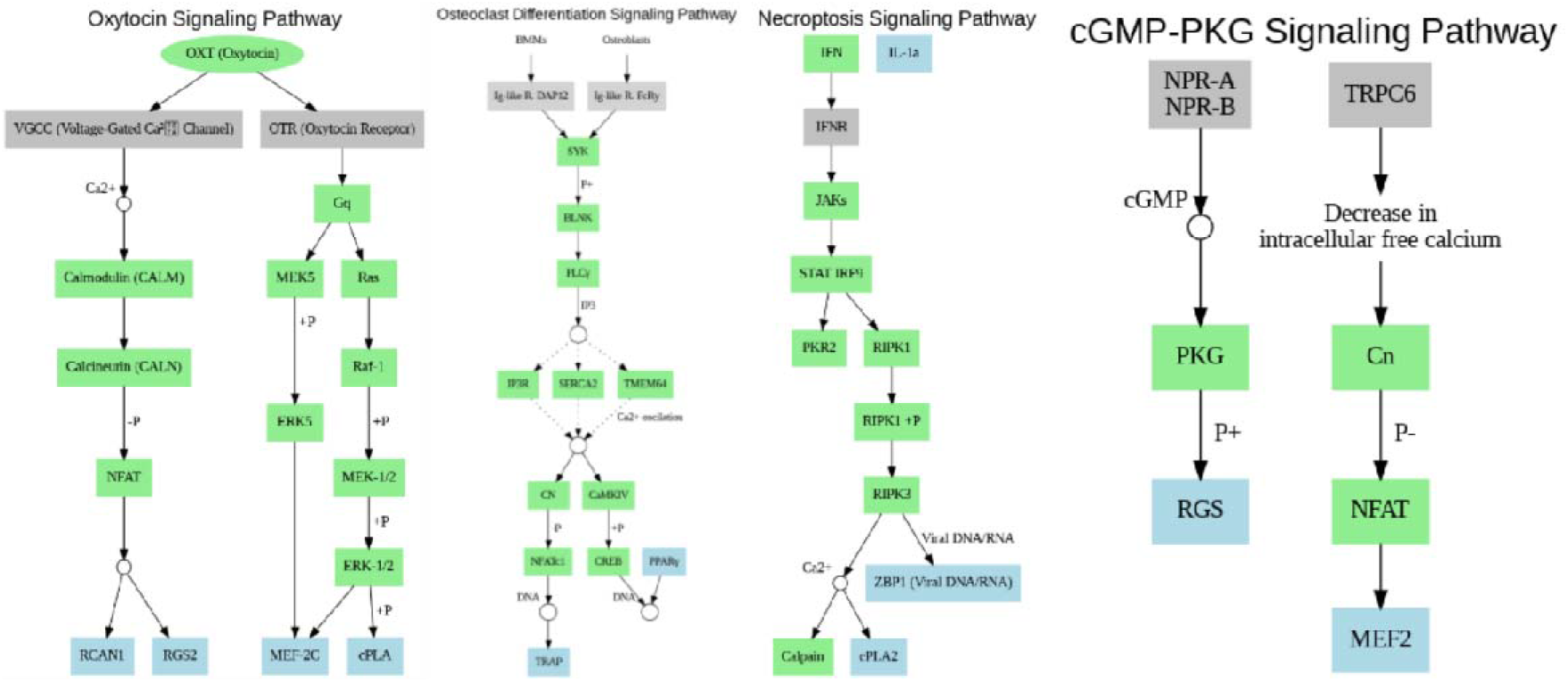
Signaling Pathways. This figure shows four important immune related pathways and explains how they help control immune responses and cell activities. The gray boxes represent cell membrane, the green boxes represent genes/proteins, and the blue boxes represent significant/immune genes. The solid line arrows are indicating direct interactions, whereas the dotted line arrows indicate indirect or regulatory interactions. Image made using python code

From the upregulated genes, five immune-related genes were identified: RCAN1, TIMD4, PPARG, SPAG5, and TLR10. From the downregulated genes, three immune-related genes were found: ZBP1, RGS2, and IL1A. Each gene was assessed for its presence in two reference sources: innate immune genes were obtained from the InnateDB.

In cattle, IL1A functions as a key pro-inflammatory cytokine, mediating immune signaling and regulating prostaglandin and progesterone secretion during infection and reproduction (Gabler, C., et al. 2010, Yamanaka et al., 2021). RGS2 and RCAN1 modulate immune cell signaling and cytokine production through G-protein and calcineurin pathways, respectively (Nunn et al., 2010, Sun et al., 2011). ZBP1 detects viral DNA and triggers antiviral responses (Maelfait et al., 2017), while TIMD4 helps in clearing apoptotic cells and maintaining immune tolerance (Miyanishi et al., 2007). PPARG plays an anti-inflammatory role by regulating macrophage activity and lipid metabolism (Lefterova et al., 2014). SPAG5 may support immune cell proliferation during infection (Zhu et al., 2021). TLR10, a pattern recognition receptor, likely contributes to early detection of pathogens and activation of innate immunity in cattle (Oosting et al., 2014). This breakdown provides a clearer understanding of the immune-related transcriptional response.

### 2.2. Signaling Pathway of immune related genes

The signaling pathway analysis highlights the four biological pathways identified in the study, where immune related genes are playing a role. They are Oxytocin, Osteoclast Differentiation, Necroptosis, and cGMP–PKG. The Oxytocin signaling pathway modulates cellular responses through calcium channel regulation and downstream activation of MAPK signaling cascades, contributing to neuropeptide signaling and immune modulation (Gimpl & Fahrenholz, 2001). In the Osteoclast Differentiation pathway, immune receptors trigger intracellular calcium signaling and activate key transcription factors such as NFATc1 and CREB, essential for immune signaling integration (Takayanagi, 2007).The Necroptosis pathway, known for its role in programmed cell death, involves effector molecules such as RIPK1 and RIPK3 and is driven by calcium signaling, linking cell death to immune activation (Pasparakis & Vandenabeele, 2015). The cGMP–PKG signaling pathway further contributes by regulating intracellular calcium levels through PKG, thus connecting calcium dynamics with pathways influencing inflammation, apoptosis, and cellular homeostasis (Lucas et al., 2000). These interconnected signaling cascades emphasize calcium signaling as a central regulator across neuroendocrine, immune, and cell death processes (Berridge, 2005).

## 3. Materials and methods

### 3.1. Data Retrieval

A total of four gene expression datasets were retrieved using keywords “Expression profiling by high throughput sequencing” AND “Immune response” from the NCBI Gene Expression Omnibus (GEO) database using the accession numbers GSE45439 (PRJNA194043), GSE62048 (PRJNA263027), GSE125964 (PRJNA518101), and GSE247921 (PRJNA1040910). These datasets represent RNA-Seq data from *Bos taurus* samples under various conditions. For GSE45439, there are 14 samples (seven *Mycobacterium bovis* infected and seven control uninfected). For GSE62048, there are 35 samples (21 control and 14 *Mycobacterium avium* subsp. paratuberculosis, (MAP) Infected). For GSE125964, there are 30 samples (ten healthy, ten clinical and ten sub clinical postpartum uterine disease). For GSE247921, there are 16 samples (eight *Mycobacterium avium* subsp. paratuberculosis, (MAP) infection and eight no treatment).

### 3.2. Sequence Data Processing

Raw RNA sequencing reads in FASTQ format were downloaded using the NCBI SRA Toolkit. The quality of raw reads was assessed using FastQC (Andrews, 2010). Low-quality bases and adapter sequences were trimmed using TrimmomaticSE (for Single ends), TrimmomaticPE (for Paired ends) (Bolger et al., 2014) parameters are ILLUMINACLIP:2:30:10:2:True was used to remove adapter sequences; LEADING:25 and TRAILING:25 were applied to trim low-quality bases from the start and end of reads; SLIDINGWINDOW:4:15 trimmed reads when the average quality in a 4-base window fell below 15; MINLEN:36 discarded reads shorter than 36 bases; AVGQUAL:25 filtered out reads with an average quality below 25; and MAXINFO:36:0.5 was used to balance read length and quality. and post-trimming quality was again evaluated using FastQC to ensure improvements in read quality.

### 3.3. Read Alignment and Post-processing

High-quality trimmed reads were aligned to the *Bos taurus* reference genome GCF_002263795.3 using Bowtie2 with default parameters. The resulting SAM files were converted to the BAM format and processed using Samtools (Li et al., 2009). This includes sorting, indexing, fixing-mate information, marking duplicates, generating alignment statistics for each sample and converting into a final BAM format.

### 3.4. Quantification of feature counts

Transcript-level quantification was carried out using FeatureCounts (Liao et al., 2014), a widely used read summarization tool. Aligned reads were assigned to genomic features based on the *Bos taurus* GTF annotation file, and raw count matrices were generated for each dataset. Strand-specific and multi-mapping read options were applied to improve the accuracy of feature assignment. The resulting count data were then used for downstream differential expression analysis, providing a comprehensive overview of gene expression across various experimental conditions.

### 3.5. Differential Gene Expression Analysis

The raw count data from all four datasets were combined and normalized using the DESeq2 package in R (Love et al., 2014). Differential expression analysis was carried out to identify genes with statistically significant changes in expression between the defined conditions. Genes with an adjusted p-value < 0.05 and log2 fold change| ± 1.5 were considered significantly differentially expressed.

### 3.6. Meta-analysis

To identify genes consistently differentially expressed across all datasets, a meta-analysis was performed using the metaRNASeq package in R (Rau et al., 2014). This method integrates results from multiple studies to improve statistical power and reduce dataset-specific biases. P-values from individual differential expression analyses were combined using Fisher’s method, allowing the detection of genes that show consistent expression patterns despite variations in experimental conditions. Only genes present across all datasets were considered to ensure comparability. Although the meta-analysis focused on integrating p-values, log2 fold change values from each dataset were used to interpret the direction and magnitude of expression changes in the significant genes.

To identify immune-related genes, the list of differentially expressed genes (DEGs) obtained from the meta-analysis was cross-referenced with InnateDB database, enabling the selection of genes involved in host immune responses.

The following tools were used to perform statistical analysis, visualization, and pathway enrichment throughout this study: R (version 4.3.1) and the ggplot2 package (version 3.4.0) were used to create the volcano plot, allowing for intuitive representation of fold-change and significance levels. ShinyGO (version 0.82) (Ge et al., 2020) was employed for pathway enrichment analysis, leveraging integrated GO and KEGG databases for functional annotation.

## 4. Discussion

This meta-analysis of bovine RNA-Seq datasets offers an integrated and high confidence view of the transcriptional landscape underlying immune responses in *Bos taurus* across diverse pathological conditions. By aggregating data from four independent studies representing infections with *Mycobacterium bovis, Mycobacterium avium* subspecies *paratuberculosis*, and uterine diseases, we overcame limitations associated with individual dataset heterogeneity, small sample sizes, and platform specific noise.

Our analytical approach integrated established bioinformatics tools and statistical methods to ensure robust and reproducible results. The workflow encompassed quality assessment of raw sequencing data using FastQC, adapter trimming and quality filtering using Trimmomatic, read alignment to the bovine reference genome using Bowtie2, post-alignment processing with SAMtools, feature quantification using FeatureCounts, and differential expression analysis using DESeq2. The meta-analysis was performed using the metaRNASeq package in R, which implements Fisher’s method for combining p-values across studies while accounting for the statistical dependencies inherent in transcriptomic data

To ensure biological relevance and functional interpretation of our findings, we cross-referenced the identified differentially expressed genes with established immune gene databases, particularly the InnateDB database, which provides curated information about genes involved in innate and adaptive immune responses. Pathway enrichment analysis was conducted using KEGG (Kyoto Encyclopedia of Genes and Genomes) pathway databases to identify the biological processes and molecular mechanisms most significantly affected by the observed gene expression changes.

The analysis revealed 28 significantly differentially expressed genes (DEGs), among which 8 were immune-related and functionally annotated through InnateDB and literature evidence. Key immune effectors such as IL1A, RGS2, RCAN1, ZBP1, PPARG, TIMD4, SPAG5, and TLR10 exhibited consistent expression alterations, reinforcing their potential as conserved biomarkers of bovine immune function.

Importantly, pathway enrichment analyses identified four crucial immune-related signaling pathways—Necroptosis, Oxytocin Signaling, Osteoclast Differentiation, and cGMP–PKG Signaling—which together underscore the intricate regulation of inflammatory responses, immune cell activation, apoptosis, and neuroimmune interactions in cattle. Calcium signaling emerged as a unifying mechanism across these pathways, highlighting its central role in coordinating immune cell function and fate decisions under stress or infection.

These findings have practical value for cattle farming and veterinary medicine. The identified genes could serve as biomarkers to detect diseases early, assess immune strength, and predict treatment success. For breeding programs, these genes could help select cattle with better disease resistance. Understanding how pathogens suppress immune responses might also lead to better vaccines and treatments.

While this study provides valuable insights, it has limitations. We only had four datasets to work with, and the different experimental conditions may have affected our results. Future research should include experimental validation of these candidate genes through qRT-PCR, protein-level assays, and functional studies using knockout or overexpression models. Expanding the meta-analysis to include newer RNA-Seq datasets, single-cell transcriptomics, and epigenomic data would further refine our understanding of context- specific and cell-type-specific immune regulation in cattle.

In conclusion, this study demonstrates the power of integrative transcriptomic meta-analysis in livestock immunogenomics and provides a valuable resource for the bovine research community. By illuminating conserved immune pathways and key transcriptional regulators, it paves the way for improving cattle health, productivity, and disease resilience through informed genomic and biotechnological strategies.

## Data and Code Availability

The processed data and scripts are available at https://github.com/vennilakm04/Transcriptome-Data-analysis

## Author contributions

VKDM & KM compiled, processed, and curated the data. VKDM & KM also ran the analysis and wrote the first draft of the manuscript. RMY conceived the project, supervised, and finalized the manuscript.

## Competing Interests

All authors declare no financial or non-financial competing interests.

## Acknowledgements

The authors acknowledge SASTRA Deemed to be University for infrastructural support.

## Notes

https://github.com/vennilakm04/Transcriptome-Data-analysis

### Competing Interest Statement

The authors have declared no competing interest.

### Summary of Updates

The p-values in the table 1 had missing decimal points. Hence, made the changes and uploaded the modified manuscript. Conclusions do not change.

